# The utility of Saw-scaled viper, *Echis pyramidum*, and Kenyan sand boa, *Eryx colubrinus* as bioindicator of heavy metals bioaccumulation in relation to DPTA soil extract and biological consequences

**DOI:** 10.1101/311357

**Authors:** Doha M. M. Sleem, Mohamed A. M. Kadry, Eman M. E. Mohallal, Mohamed A. S. Marie

## Abstract

The present investigation was conducted to compare between the ecotoxocological effects of Ca, Mg, Fe, Cu, Zn, Co, Mo, Mn, B, Al, Sr, Pb, Ni, Cd and Cr on the saw-scaled viper, *Echis pyramidum (E. p.)* and the Kenyan sand boa, *Eryx colubrinus* (*E*.c.) inhabiting Gabal El-Nagar and Kahk Qibliyyah respectively in El-Faiyum desert, Egypt. Accumulation varied significantly among the liver, kidney and muscle. The relationship between concentrations of heavy metals in snakes and those in the soil from the collected sites was established by analyzing metal DPTA in soil. Bioaccumulation factor is calculated to estimate the degree of toxicity within the tissues. Morphometric analysis was recorded. All body morphometric measurements were higher in *E. p*. than in *E. c.*. Body, liver, gonad, kidney and heart weight, HSI, GSI, RBCs count, Hb content, PCV, MCV, MCH, MCHC, plasma glucose, total lipids and total proteins showed a significant increase in *E. p*. Histopathological examination showed damage and alterations of liver, kidney and testes sections. The tissues of *E. c*. were more destructed than those of *E. p.*.

## Introduction

Little is known about snakes in the field of ecotoxicology and it needs several investigations **(Albers *et al.*, 2000; Hopkins, 2000; Sparling *et al.*, 2000 and Selcer, 2005)**; because snakes are cold blooded (poikilothermic) animals, and they have low rates of catabolism, thus, they may maintain higher body burdens of contaminants **(Gibbons *et al*., 2000)**. As environmental contaminant is considered one of the major six factors, that contributes to the decline of the snake populations **(Todd *et al.*, 2010)**, therefore, studies have been increased over the last decade seeking for the conservation of the snakes, and assert on the role played by the bioindicators in the change of the response to a certain stress, and this will guide to the endangered snake species **(Hopkins *et al.*, 1999&2001; Burger *et al.*, 2005, 2006&2007; Campbell *et al.*, 2005; Albrecht *et al.*, 2007; Wylie *et al.*, 2009 and Rezaie-Atagholipour *et al.*, 2012)**.

The objective of this study was to determine whether the saw-scaled viper, *Echis pyramidum* or the Kenyan sand boa, *Eryx colubrinus* as bioindicator of the internal exposure of metal.

In this investigation the level of heavy metals in the soil were analyzed, because snakes are mainly found on the top surface of the soil, which is mainly contaminated with more heavy metals than the bottom **(Olafisoye *et al.*, 2013)**.

Since the environmental contaminants exert deleterious effects on snakes causing alteration of morphometric characteristics, genetic effects, behavioral parameters and tissue residue levels **(Irwin and Irwin, 2005 and Walker *et al.*, 2012)**, morphometrics, hematology and plasma biochemistry, and histopathology were conducted in this current work to aid in choosing the suitable bioindicator.

## Materials and methods

### Collection site

Eight specimens of both types of selected snakes *Echis pyrimidum* and *Eryx colubrinus* were collected from two different sites in El-Faiyum desert, Egypt.

#### Site 1 [Gabal El-Nagar] (Fig.1.)

Specimens of saw-scaled viper; *Echis* pyrimidum as well as soil samples were collected during the summer season from Gabal El-Nagar (Mahatet El-Rafa); a rocky area near a water drainage and a cultivated area planted with wheat.

Global Position System (GPS) of El-Faiyum: [29° 22’ N 30° 37’E].

**Fig. 1.**
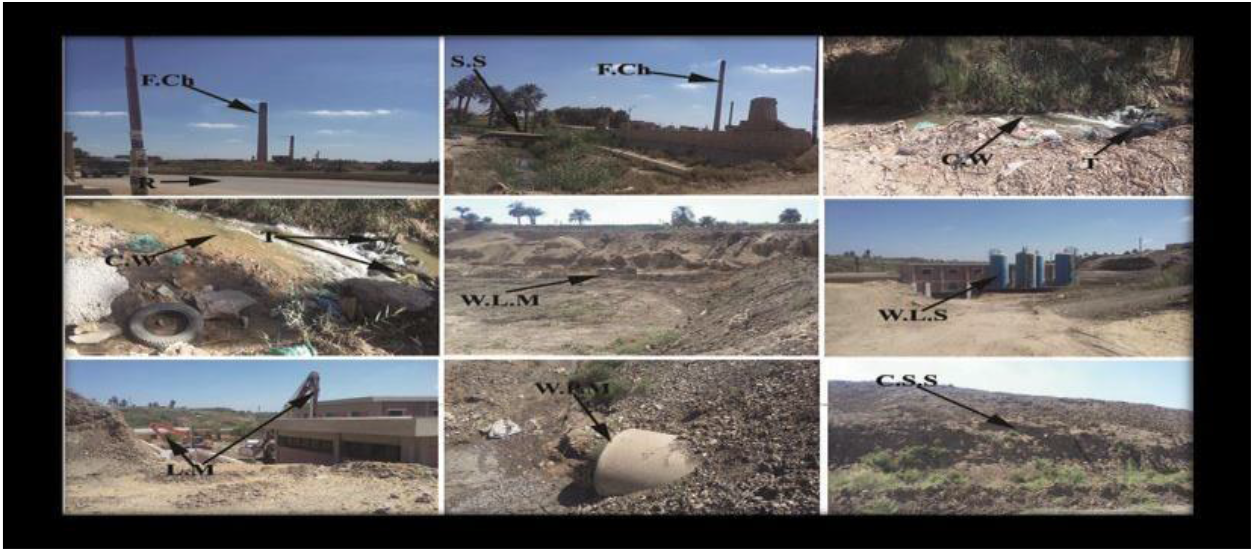
Photomicrograph shows contaminated sites in Gabal El-Nagar. C.W: Contaminated water; T: Trash; W.L.M: Water lifting machine; W.L.S: Water lifting station; L.M: Lifting machine; W.P.M: Water pumping machine C.S.S: Collected soil samples.

#### Site 2 [Kahk Qibliyyah, Abshowy] (Fig.2.)

Specimens of Kenyan sand boa; Eryx *colubrinus* as well as soil samples were collected during the summer season from Kahk Qibliyyah; a planted area near a sandy road. Global Positioning System (GPS): [29° 24’ N 30° 38’ E]

**Fig. 2.**
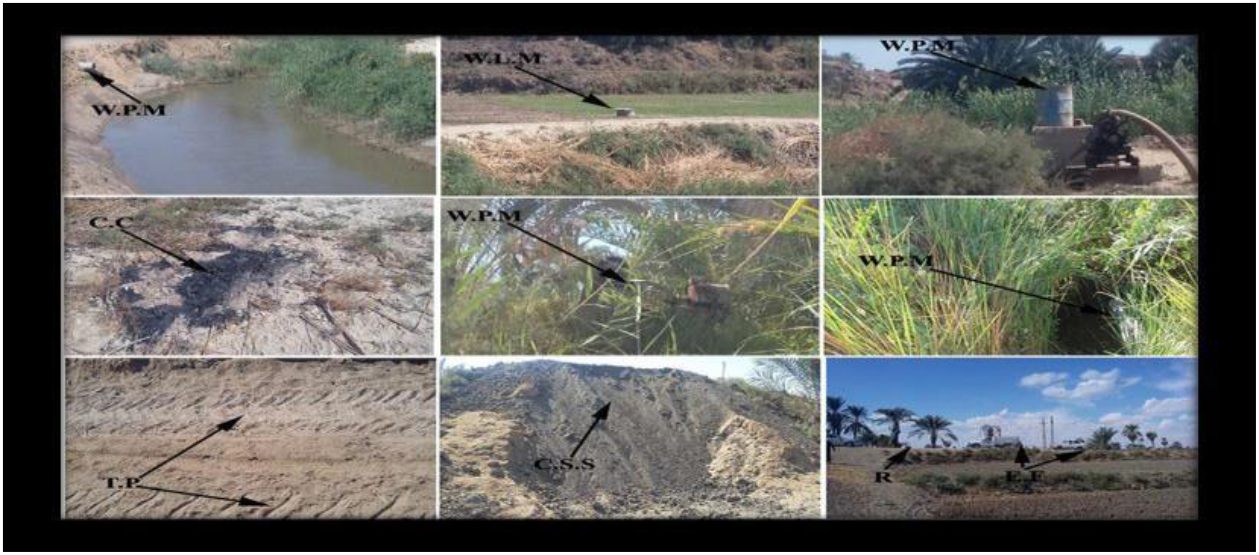
Photomicrograph shows contaminated site in Kahk Qibliyyah. C.C: Contaminated coal; T.P: Tractor’s print; C.S.S: Collected soil samples; R: Road; E.F: Exhausted fumes; W.P.M.: Water Pumping Machine and W.L.M.: Water Lifting Machine.

### Morphometric measurements

The snakes were separated and anaesthetized, SVL (Snout-vent length) (mm), TL (Tail length) (mm), TBL (Total body length) (mm), and HL (Head length) (mm) were measured by using Vernier caliper. Body, liver, heart, and kidney weight (g); hepatosomatic index (HIS), and gonadosomatic index (GSI) were estimated. These data gives a precise evaluation of the status of the snake’s body during the study **(Gómez *et al*., 2016)**.

### Blood Collection and hematology

Blood samples were taken from the heart into the heparinized eppendrofs after the snakes being decapitated. Neubauer haemocytometer was used to determine red blood corpuscles count (RBCs). Hemoglobin was estimated spectrophotometrically in whole blood collected in heparin according to the method of **Drabkin and Austin (1932)** using a kit of Biodiagnostics Company. Hematocrit centrifuge was used to measure packed cell volume (PCV). Blood indices [Mean Corpuscular Volume (MCV), Mean Corpuscular Hemoglobin (MCH), and Mean Corpuscular Hemoglobin Concentration (MCHC)] were calculated according to **Gupta (1970)**.

### Plasma Biochemistry

A part of blood was centrifuged for five minutes at 10000 rpm. The plasma was removed, placed in another clean eppendorf tubes, immediately frozen at −20oC and stored for biochemical analyses. Plasma glucose was estimated spectrophotometrically according to the method of **Trinder (1969)** using a kit of Biodiagnostics Company. Plasma total lipids were estimated spectrophotometrically according to the method of **Zöllner and Kirsch (1962)** using a kit of Biodiagnostics Company. Plasma total proteins were determined spectrophotometrically according to the method of **Gornall *et al*. (1949)** using the kit of Biodiagnostics company.

### Heavy metals bioaccumulation in liver, kidney, and muscle tissues of snakes

0.5 g of each tissue, 7 ml of HNO3 (65%) and 1 ml of H2O2 (30%) reagents were added in a closed vessel to be put inside the temperature control microwave digestion for the metal determination by the spectroscopic method. The samples were left to be cooled for 24 hours in the room temperature, and then the digested solutions were transferred to 25 ml conical to be diluted; finally, they were ready to be measured by using ICAP (Inductive Coupled Aragon Plasma), after they were calibrated at 0.05 pp^m^.

### Heavy metals in available soil

The determination of heavy metals in available soil was according to the method of **Soltanpour and Schwab (1977)**.

**Bioaccumulation Factor (BAF)**

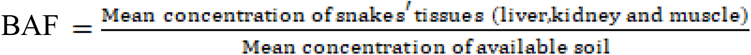

### Histopathological examination

Autopsy samples were taken from the liver, kidney and testes of the two selected studied snakes according to **(Bancroft and Gamble, 2002)**.

### Statistical analysis

Pairwise significance tests were made using Student“s ***t-test*** to compare between Kenyan sand boa, *Eryx colubrinus* and saw-scaled viper, *Echis pyrimidum* in all studied blood parameters and heavy metals bioaccumulation in their different tissues. A probability level at **(P<0.05)** was considered as significant, at **(P<0.01)** was considered as highly significant and at **(P<0.001)** was considered as very highly significant.

All statistics were carried out using statistical analysis program; Predictive Analytics Software **(PASW statistics 18.0 Release 18.0.0)**.

### Ethical considerations

We follow scientific ethics in selection and handling of snakes. Our Institutional Animal Care and Use Committee (IACUC) at Zoology Department, Faculty of Science, Cairo University has approved this study protocol from the ethical point of view and according to Animal welfare Act of the Ministry of Agriculture in Egypt that enforces the humane treatment of animal.

## Results

Body morphometric analyses showed that the mean values of all studied measured parameters in the saw-scaled viper, *Echis pyramidum* (*E. p*.) were higher than those in the Kenyan sand boa, *Eryx colubrinus* (*E. c*.) (Table 1).

**Table (1):**
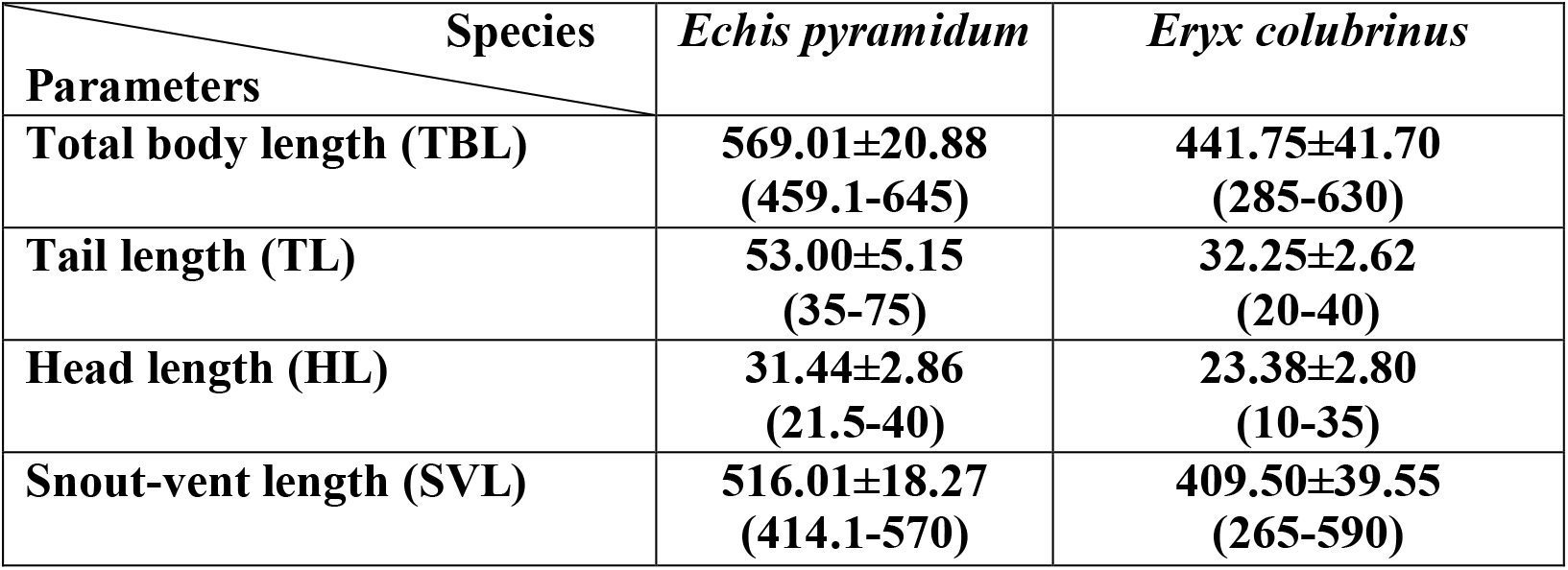
Morphometric measurements (mm) of the saw-scaled viper, *Echis pyramidum* and the Kenyan sand boa, *Eryx colubrinus* inhabiting El-Faiyum desert, Egypt.

Changes in body weight, liver, gonads, kidney and heart weight; hepato-somatic index (HSI) and gonado-somatic index (GSI) were higher in the saw-scaled viper, *Echis pyramidum* (*E. p*.) than in the Kenyan sand boa, *Eryx colubrinus* (*E. c*.), and showed a very highly significant increase (P < 0.001) (Table 2).

**Table (2):** Changes in body weight, liver, gonad, kidney and heart weight (g); hepato-somatic index (HSI) and gonado-somatic index (GSI) between saw-scaled viper, *Echis pyramidum* and Kenyan sand boa, *Eryx colubrinus* inhabiting El-Faiyum, Egypt.

| Parameters \ Species | *Echis pyramidum* | *Eryx colubrinus* | *t*-values |
| --- | --- | --- | --- |
| Body weight (g) | 94.11±3.24 | 86.91±26.80 | 6.91*** |
| Liver weight (g) | 4.48±0.47 | 1.80±0.35 | 7.03*** |
| Hepato-somatic index (HSI) $\times 10^{-4}$ | 52.78±2.09 | 28.13±4.95 | 9.85*** |
| Gonad weight (g) | 1.42±0.33 | 1.33±0.45 | 5.09*** |
| Gonado-somatic index (GSI) $\times 10^{-5}$ | 4.68±0.51 | 1.53±0.38 | 4.25*** |
| Kidney weight (g) | 1.49±0.10 | 0.69±0.11 | 8.62*** |
| Heart weight (g) | 0.71±0.06 | 0.33±0.06 | 8.22*** |

Data are represented as means of eight (8) samples ± S.E.
*t*-values = Student *t*-test between *Echis pyramidum* and *Eryx colubrinus* for each studied parameter.
\*\*\* Very highly significant difference at ( $P < 0.001$ ).

*Echis pyramidum* (*E. p*.) showed a very highly significant increase (P < 0.001) in all studied blood parameters; Red blood corpuscles (RBCs) count, Hemoglobin content (Hb) and Packed cell volume (PCV) than in *Eryx colubrinus* (*E. c*.) (Table 3).

**Table (3):** Blood parameters of saw-scaled viper, *Echis pyramidum* and Kenyan sand boa, *Eryx colubrinus* inhabiting El-Faiyum desert, Egypt.

| **Species**<br>**Parameters** | *Echis pyramidum* | *Eryx colubrinus* | *t-values* |
| --- | --- | --- | --- |
| **RBCs ($\times 10^6$ /mm<sup>3</sup>)** | **1.43±0.14** | **0.69±0.14** | **7.82\*\*\*** |
| **Hb (g/100 ml)** | **33.61±4.55** | **7.27±1.07** | **5.01\*\*\*** |
| **PCV (%)** | **28.26±1.13** | **27.88±5.14** | **11.04\*\*\*** |

Data are represented as means of eight samples ± S.E.
*t-values* = Student *t-test* between *Echis pyramidum* & *Eryx colubrinus* for each estimated blood parameters.
\*\*\* Very highly significant difference ( $P < 0.001$ )

*Echis pyramidum* (*E. p*.) showed a very highly significant increase (P < 0.001) in calculated blood indices; mean corpuscular volume (MCV), mean corpuscular hemoglobin (MCH) and mean corpuscular hemoglobin concentration (MCHC) than in *Eryx colubrinus* (*E. c*.) (Table 4).

**Table (4):** Calculated blood indices of saw-scaled viper, *Echis pyramidum* and Kenyan sand boa, *Eryx colubrinus* inhabiting El-Faiyum desert, Egypt.

| **Species**<br>**Parameters** | *Echis pyramidum* | *Eryx colubrinus* | *t-values* |
| --- | --- | --- | --- |
| **MCV ($\times 10^{-4}$ fl)** | **6.79±0.72** | **4.79±0.58** | **11.22\*\*\*** |
| **MCH ($\times 10^{-4}$ pg/cell)** | **6.18±0.52** | **3.66±0.99** | **7.79\*\*\*** |
| **MCHC (g/100 ml)** | **44.20±2.57** | **34.46±8.62** | **8.69\*\*\*** |

Data are represented as means of eight samples ± S.E.
*t-values* = Student *t-test* between *Echis pyramidum* & *Eryx colubrinus* for each estimated blood parameters.
\*\*\* Very highly significant difference ( $P < 0.001$ ).

*Echis pyramidum* (*E. p*.) recorded a very highly significant increase (P < 0.001) in plasma glucose, total lipids and total protein than in *Eryx colubrinus* (*E. c*.) (Table 5).

**Table (5):** Changes in plasma glucose, total lipids and total protein of saw-scaled viper, *Echis pyramidum* and Kenyan sand boa, *Eryx colubrinus* inhabiting El-Faiyum desert, Egypt.

| **Parameters \ Species** | ***Echis pyramidum*** | ***Eryx colubrinus*** | ***t-values*** |
| --- | --- | --- | --- |
| **Glucose (mg/100 ml)** | **89.91±1.75** | **56.60±14.32** | **8.95\*\*\*** |
| **Total lipids (mg/100 ml)** | **7293.75±424.18** | **1263.15±363.57** | **5.19\*\*\*** |
| **Total protein (g/100 ml)** | **6.36±0.22** | **3.26±0.68** | **9.07\*\*\*** |

Data are represented as means of eight samples ± S.E.
***t-values*** = Student *t-test* between *Echis pyramidum* & *Eryx colubrinus* for each estimated blood parameters.
\*\*\* Very highly significant difference ( $P < 0.001$ )

Essential heavy metals [Ca, Fe, Cu, Zn] in soils collected from Kahk Qibliyyah showed a highly significant increase (P < 0.001) than those in soils collected from Gabal El-Nagar, except Mo, which was significantly higher in Gabal El Nagar [0.085±0.005] than in Kahk Qibliyyah [0.038±0.009] (P < 0.001). Mg concentrations recorded a highly significant increase (P < 0.01) in soils collected from Gabal El-Nagar compared to those collected from Kahk Qibliyyah. Co and Mn concentrations revealed a significant increase (P < 0.05) in soils collected from Kahk Qibliyyah than those collected from Gabal El-Nagar (Table 6).

**Table (6):** Concentration of essential heavy metals (ppm) in the available soils collected from Gabal El-Nagar and Kahk Qibliyyah in El-Faiyum desert, Egypt.

| Soil samples<br>Heavy metals | El-Faiyum Desert |  | *t*-values |
| --- | --- | --- | --- |
|  | *Site I*<br>Gabal El-Nagar | *Site II*<br>Kahk Qibliyyah |  |
| Calcium (Ca) | 39.418±7.047<br>(9.11-58.68) | 52.547±5.342<br>(35.98-63.7) | 9.873*** |
| Magnesium (Mg) | 650.115±269.496<br>(62.66-1913.6) | 509.267±57.606<br>(328.6-619.8) | 4.356** |
| Iron (Fe) | 9.333±2.702<br>(1.817-21.24) | 21.747±2.868<br>(15.928-30.68) | 5.861*** |
| Copper (Cu) | 1.582±0.375<br>(0.486-3.262) | 5.137±0.475<br>(3.68-6.176) | 5.521*** |
| Zinc (Zn) | 5.043±0.482<br>(3.756-6.958) | 6.292±0.66<br>(4.682-8.248) | 13.093*** |
| Cobalt (Co) | 0.159±0.02<br>(0.066-0.205) | 0.733±0.132<br>(0.347-1.064) | 4.149* |
| Molybdenum (Mo) | 0.085±0.005<br>(0.067-0.094) | 0.038±0.009<br>(0.016-0.067) | 6.919*** |
| Manganese (Mn) | 10.101±2.44<br>(4.956-21.3) | 68.593±12.092<br>(40.4-105.06) | 3.712* |

Data are represented as means of six samples ± S.E. Lower and upper values between parentheses.
*t*-values = Student *t*-test between the studied sites in El-Faiyum desert for each estimated heavy metal.
\* Significant difference ( $P < 0.05$ ), \*\* Highly significant difference ( $P < 0.01$ ), \*\*\* Very highly significant difference ( $P < 0.001$ ).

Non-essential heavy metals [Pb & Cr] collected from Kahk Qibliyyah showed a highly significant increase (P < 0.001) than those in soils collected from Gabal El-Nagar except Sr which was significantly higher in Gabal El-Nagar [0.708±0.135] than in Kahk Qibliyyah [0.698±0.082] (P < 0.001). B concentration recorded a highly significant increase (P < 0.01) in soils collected from Gabal El-Nagar compared to those collected from Kahk Qibliyyah, while Cd concentration showed that a highly significant increase (P < 0.01) in soils collected from Kahk Qibliyyah compared to those collected from Gabal El-Nagar. Ni concentration revealed a significant increase (P < 0.05) in soils collected from Kahk Qibliyyah than those in soils collected from Gabal El-Nagar. Al concentration was higher in Gabal El-Nagar than in Kahk Qibliyyah and showed insignificant difference (Table 7).

**Table (7):** Concentration of non-essential heavy metals (ppm) in the available soils collected from Gabal El-Nagar and Kahk Qibliyyah in El-Faiyum desert, Egypt.

| Species<br>Heavy metals | El-Faiyum Desert |  | *t*-values |
| --- | --- | --- | --- |
|  | *Site I*<br>Gabal El-Nagar | *Site II*<br>Kahk Qibliyyah |  |
| Boron (B) | 2.571±0.774<br>(0.749-6.06) | 1.616±0.51<br>(0.559-3.2) | 4.505** |
| Aluminium (Al) | 1.431±0.505<br>(0.048-2.814) | 0.058±0.003<br>(0.048-0.064) | 2.345 |
| Strontium (Sr) | 0.708±0.135<br>(0.266-1.129) | 0.698±0.082<br>(0.485-0.932) | 9.347*** |
| Lead (Pb) | 0.223±0.048<br>(0.048-0.383) | 0.824±0.04<br>(0.696-0.893) | 5.485*** |
| Nickel (Ni) | 0.186±0.05<br>(0.027-0.384) | 0.836±0.16<br>(0.452-1.313) | 4.042* |
| Cadmium (Cd) | 0.031±0.006<br>(0.015-0.049) | 0.053±0.016<br>(0.028-0.104) | 4.809** |
| Chromium (Cr) | 0.05±0.004<br>(0.043-0.07) | 0.064±0.004<br>(0.05-0.071) | 16.08*** |

Data are represented as means of six samples ± S.E. Lower and upper values between parentheses.
*t*-values = Student *t*-test between the studied sites in El-Faiyum desert for each estimated heavy metal.
\* Significant difference ( $P < 0.05$ ), \*\* Highly significant difference ( $P < 0.01$ ), \*\*\* Very highly significant difference ( $P < 0.001$ ).

The Saw-scaled viper, *Echis pyramidum* bioaccumulated a very highly significant concentrations (P < 0.001) of some essential heavy metals (Mg, Zn and Mn) in their studied tissues (liver, kidney and muscle) than in the Kenyan sand boa, *Eryx colubrinus*. Fe was bioaccumulated in the Kenyan sand boa, *Eryx colubrinus* in very highly significant concentartions in liver (P < 0.001), and highly significant concentrations in kidney (P < 0.01), and it was higher in muscle with insignificant difference compared to the saw-scaled viper, *Echis pyramidum*. While Mo was highly significant bioaccumulated (P < 0.01) in liver and kidney tissues of the Kenyan sand boa, *Eryx colubrinus*, it revealed a significant increase (P < 0.05) in muscle of the Kenyan sand boa, *Eryx colubrinus* than in the saw-scaled viper, *Echis pyramidum*. The bioaccumulation of Ca in liver tissue showed a very highly significant increase (P < 0.001) in the Kenyan sand boa, *Eryx colubrinus* than in the saw-scaled viper, *Echis pyramidum*, while kidney and muscle tissues of the saw-scaled viper, *Echis pyramidum* recorded a very highly significant increase (P < 0.001) than in the Kenayn sand boa, *Eryx colubrinus*. The bioaccumulation of Cu in kidney and muscle tissues of the Kenayn sand boa, *Eryx colubrinus* were very highly significant increase (P < 0.001) and highly significant increase (P < 0.01) respectively but it recorded a very highly significant increase (P < 0.001) in liver of the saw-scaled viper, *Echis pyramidum* than in the Kenyan sand boa, *Eryx colubrinus*. Co was bioaccumulated in liver and kidney tissue of the Kenayn sand boa, *Eryx colubrinus* than in the saw-scaled viper, *Echis pyramidum* with very highly significant difference (P < 0.001) while in muscle tissues, Co was detected only in the Kenyan sand boa, *Eryx colubrinus* (Table 8).

**Table (8):**
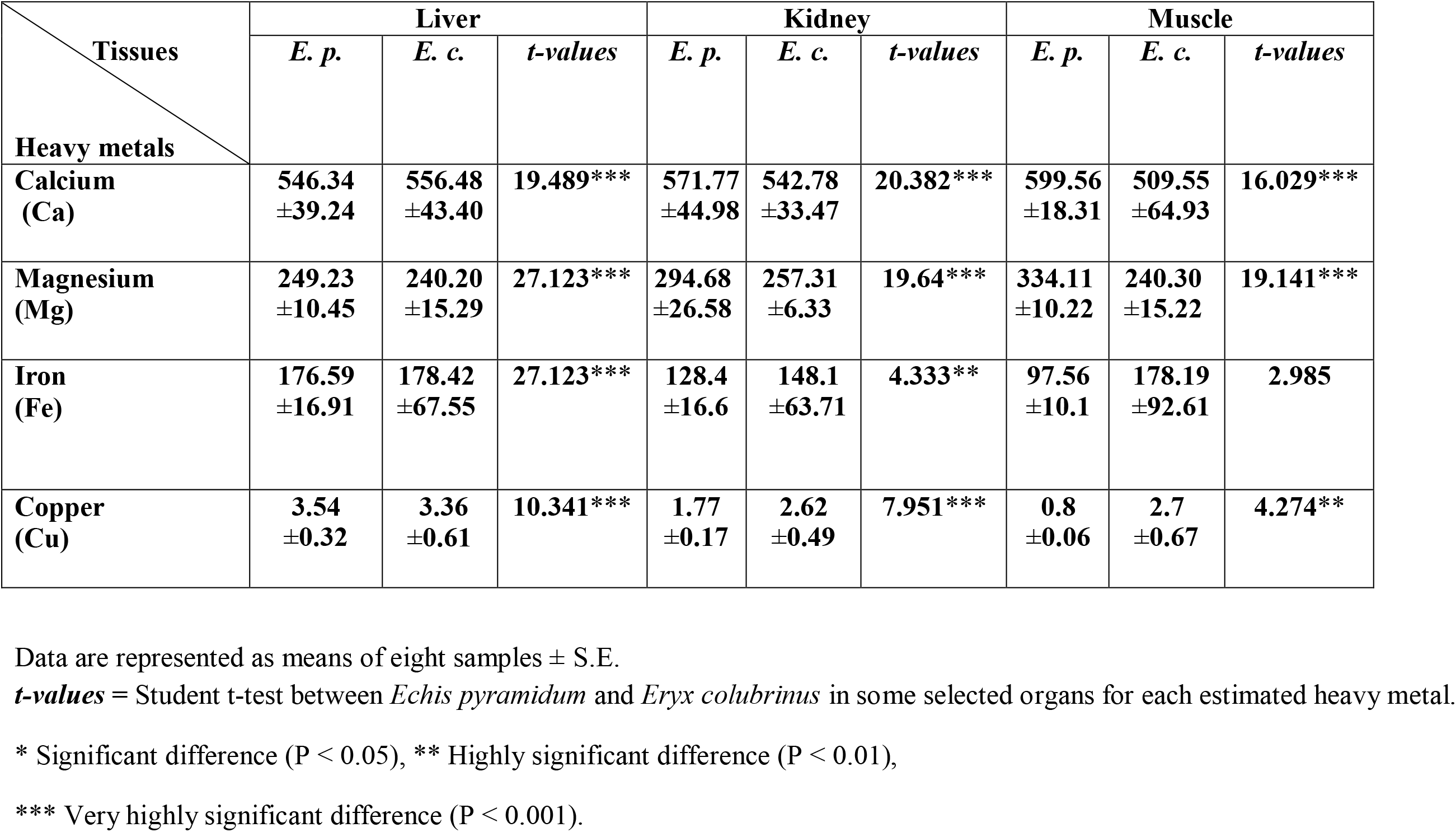
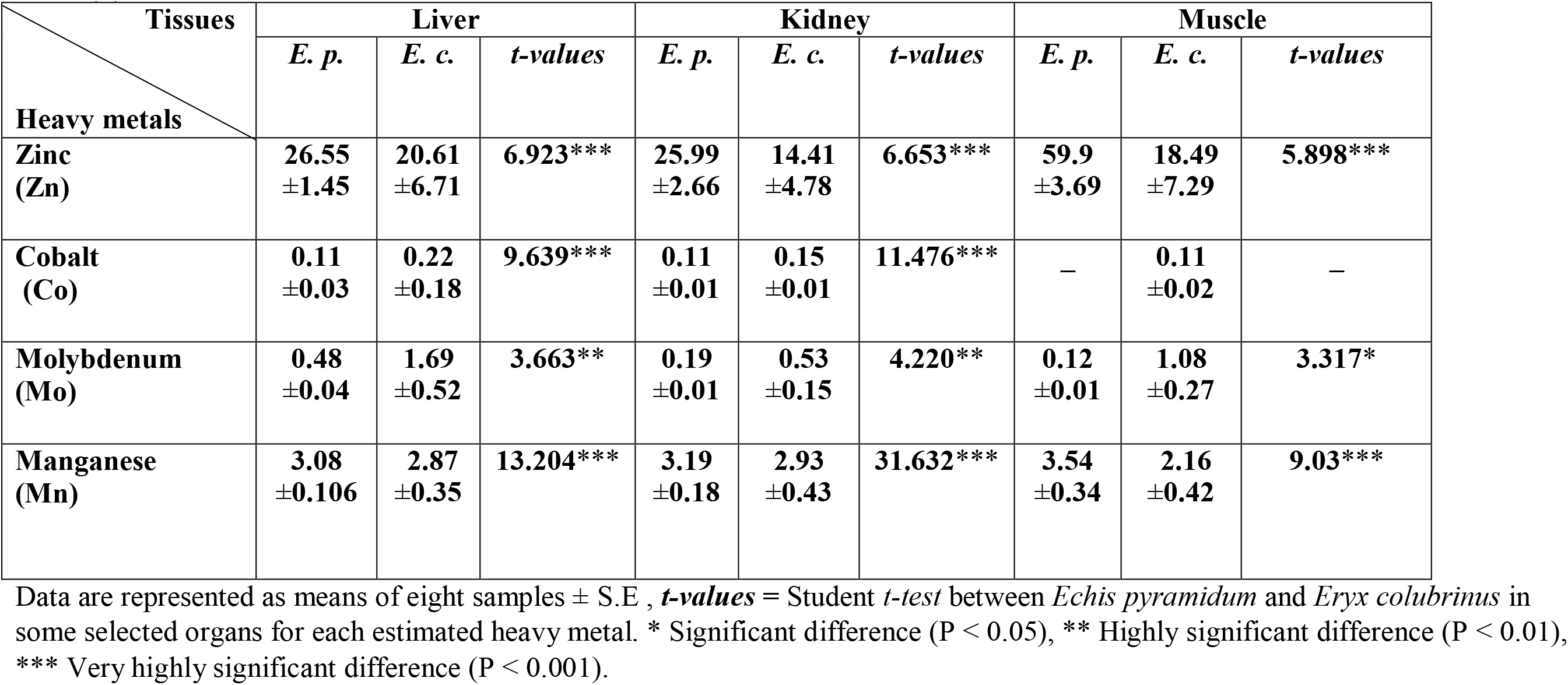
Bioaccumulation of essential heavy metals (Ca, Mg, Cu, Fe, Zn, Co, Mo and Mn) in liver, kidney and muscle (ppm) of saw-scaled viper, *Echis pyramidum* (*E. p*.) and Kenyan sand boa, *Eryx colubrinus* (*E. c*.) inhabiting El-Faiyum desert, Egypt.

| Tissues<br>Heavy metals | Liver |  |  | Kidney |  |  | Muscle |  |  |
| --- | --- | --- | --- | --- | --- | --- | --- | --- | --- |
|  | *E. p.* | *E. c.* | *t-values* | *E. p.* | *E. c.* | *t-values* | *E. p.* | *E. c.* | *t-values* |
| Calcium (Ca) | 546.34<br>±39.24 | 556.48<br>±43.40 | 19.489*** | 571.77<br>±44.98 | 542.78<br>±33.47 | 20.382*** | 599.56<br>±18.31 | 509.55<br>±64.93 | 16.029*** |
| Magnesium (Mg) | 249.23<br>±10.45 | 240.20<br>±15.29 | 27.123*** | 294.68<br>±26.58 | 257.31<br>±6.33 | 19.64*** | 334.11<br>±10.22 | 240.30<br>±15.22 | 19.141*** |
| Iron (Fe) | 176.59<br>±16.91 | 178.42<br>±67.55 | 27.123*** | 128.4<br>±16.6 | 148.1<br>±63.71 | 4.333** | 97.56<br>±10.1 | 178.19<br>±92.61 | 2.985 |
| Copper (Cu) | 3.54<br>±0.32 | 3.36<br>±0.61 | 10.341*** | 1.77<br>±0.17 | 2.62<br>±0.49 | 7.951*** | 0.8<br>±0.06 | 2.7<br>±0.67 | 4.274** |
|  | *E. p.* | *E. c.* | *t-values* | *E. p.* | *E. c.* | *t-values* | *E. p.* | *E. c.* | *t-values* |
| **Zinc**<br>**(Zn)** | **26.55**<br>**±1.45** | **20.61**<br>**±6.71** | **6.923\*\*\*** | **25.99**<br>**±2.66** | **14.41**<br>**±4.78** | **6.653\*\*\*** | **59.9**<br>**±3.69** | **18.49**<br>**±7.29** | **5.898\*\*\*** |
| **Cobalt**<br>**(Co)** | **0.11**<br>**±0.03** | **0.22**<br>**±0.18** | **9.639\*\*\*** | **0.11**<br>**±0.01** | **0.15**<br>**±0.01** | **11.476\*\*\*** | – | **0.11**<br>**±0.02** | – |
| **Molybdenum**<br>**(Mo)** | **0.48**<br>**±0.04** | **1.69**<br>**±0.52** | **3.663\*\*** | **0.19**<br>**±0.01** | **0.53**<br>**±0.15** | **4.220\*\*** | **0.12**<br>**±0.01** | **1.08**<br>**±0.27** | **3.317\*** |
| **Manganese**<br>**(Mn)** | **3.08**<br>**±0.106** | **2.87**<br>**±0.35** | **13.204\*\*\*** | **3.19**<br>**±0.18** | **2.93**<br>**±0.43** | **31.632\*\*\*** | **3.54**<br>**±0.34** | **2.16**<br>**±0.42** | **9.03\*\*\*** |

Data are represented as means of eight samples ± S.E , *t-values* = Student *t-test* between *Echis pyramidum* and *Eryx colubrinus* in some selected organs for each estimated heavy metal. \* Significant difference ( $P < 0.05$ ), \*\* Highly significant difference ( $P < 0.01$ ), \*\*\* Very highly significant difference ( $P < 0.001$ ).

The present data showed that the saw-scaled viper, *Echis pyramidum* bioaccumulated a very highly significant concentrations (P < 0.001) of Sr in the selected tissues (liver, kidney and muscle) than in the Kenyan sand boa, *Eryx colubrinus*. The bioaccumulation of Cd in liver and kidney showed a very highly significant concentrations (P < 0.001) in the Kenyan sand boa, *Eryx colubrinus* than in the saw-scaled viper, *Echis pyramidum* while muscle recorded a very highly significant concentrations (P < 0.001) in the saw-scaled viper, *Echis pyramidum* than in the Kenyan sand boa, *Eryx colubrinus*. Cr bioaccumulated a very highly significantly concentrations (P < 0.001) in kidney and muscle of the Kenyan sand boa, *Eryx colubrinus* while liver of the saw-scaled viper, *Echis pyramdium* revealed a very highly significantly concentrations (P < 0.001) than the saw-scaled viper, *Echis pyramidum*. Al was very highly significantly bioaccumulated (P < 0.001) in liver and muscle of the saw-scaled viper, *Echis pyramidum* and a highly significant concentration (P < 0.01) was recorded in kidney of the saw-scaled viper, *Echis pyramidum* than in the Kenyan sand boa, *Eryx colubrinus*. B bioaccumulated a very highly significantly concentrations (P < 0.001) and highly significant concentrations

(P < 0.01) in [liver and muscle] and kidney respectively of the Kenyan sand boa, *Eryx colubrinus* than in the saw-scaled viper, *Echis pyramidum*. Ni recorded a very highly significant concentrations (P < 0.001) and significant concentrations (P < 0.05) in liver and [kidney and muscle] respectively of the saw-scaled viper, *Echis pyramidum*than in the Kenyan sand boa, *Eryx colubrinus*. Pb bioaccumulated in kidney of the saw-scaled viper, *Echis pyramidum* with a very highly significant concentrations (P < 0.001), it revealed a significant increase (P < 0.05) in liver and muscle of the Kenyan sand boa, *Eryx colubrinus* and the saw-scaled viper, *Echis pyramidum* respectively (Table 9).

**Table (9):** Bioaccumulation of non-essential heavy metals (B, Al, Sr, Pb, Ni, Cd and Cr) in liver, kidney and muscle (ppm) of saw-scaled viper, *Echis pyramidum* (*E. p*.) and Kenyan sand boa, *Eryx colubrinus* (*E. c*.) inhabiting El-Faiyum, Egypt.

| Tissues<br>Heavy<br>metals | Liver |  |  | Kidney |  |  | Muscle |  |  |
| --- | --- | --- | --- | --- | --- | --- | --- | --- | --- |
|  | *E. p.* | *E. c.* | *t-values* | *E. p.* | *E. c.* | *t-values* | *E. p.* | *E. c.* | *t-values* |
| **Boron**<br>**(B)** | **8.77**<br>**±0.24** | **19.19**<br>**±3.58** | **6.372\*\*\*** | **17.08**<br>**±3.11** | **22.83**<br>**±15.55** | **2.592\*** | **5.49**<br>**±0.23** | **10.72**<br>**±2.84** | **5.282\*\*\*** |
| **Aluminium**<br>**(Al)** | **108.75**<br>**±8.34** | **48.06**<br>**±9.00** | **7.98\*\*\*** | **72.03**<br>**±21.18** | **62.90**<br>**±26.24** | **4.132\*\*** | **132.8**<br>**±17.94** | **44.25**<br>**±9.87** | **5.856\*\*\*** |
| **Strontium**<br>**(Sr)** | **2.22**<br>**±0.13** | **0.96**<br>**±0.35** | **6.567\*\*\*** | **2.26**<br>**±0.2** | **1.03**<br>**±0.32** | **6.839\*\*\*** | **2.08**<br>**±0.81** | **1.11**<br>**±0.36** | **7.358\*\*\*** |
| **Lead**<br>**(Pb)** | **0.69**<br>**±0.082** | **2.04**<br>**±0.63** | **3.871\*** | **2.01**<br>**±0.501** | **0.95±**<br>**0.28** | **4.768\*\*\*** | **7.81**<br>**±3.85** | **0.70**<br>**±0.12** | **2.047\*** |

| Tissues<br>Heavy Metals | Liver |  |  | Kidney |  |  | Muscle |  |  |
| --- | --- | --- | --- | --- | --- | --- | --- | --- | --- |
|  | *E. p.* | *E. c.* | *t-values* | *E. p.* | *E. c.* | *t-values* | *E. p.* | *E. c.* | *t-values* |
| Nickel (Ni) | 1.70<br>±0.48 | 1.13<br>±0.41 | 4.525*** | 6.93<br>±1.72 | 1.51<br>±0.71 | 3.7* | 3.16<br>±1.06 | 1.13<br>±0.41 | 3.52* |
| Cadmium (Cd) | 0.11<br>±0.01 | 0.13<br>±0.02 | 12.778*** | 0.10<br>±0.01 | 0.12<br>±0.01 | 15.951*** | 0.11<br>±0.01 | 0.10<br>±0.01 | 18.085*** |
| Chromium (Cr) | 3.91<br>±0.31 | 2.54<br>±0.15 | 13.239*** | 1.68<br>±0.18 | 1.89<br>±0.22 | 12.556*** | 1.99<br>±0.33 | 2.49<br>±0.18 | 11.677*** |

Data are represented as means of eight samples ± S.E.
*t-values* = Student *t-test* between *Echis pyramidum* and *Eryx colubrinus* in some selected organs for each estimated heavy metal.
\* Significant difference ( $P < 0.05$ ), \*\*\* Very highly significant difference ( $P < 0.001$ ).

Bioaccumulation factor in Gabal El-Nagar was higher than in Kahk Qibliyyah except Mg and Mo, bioaccumulation factor of Co in muscle in Gabal El-Nagar was not detected (Table 10).

**Table (10):**
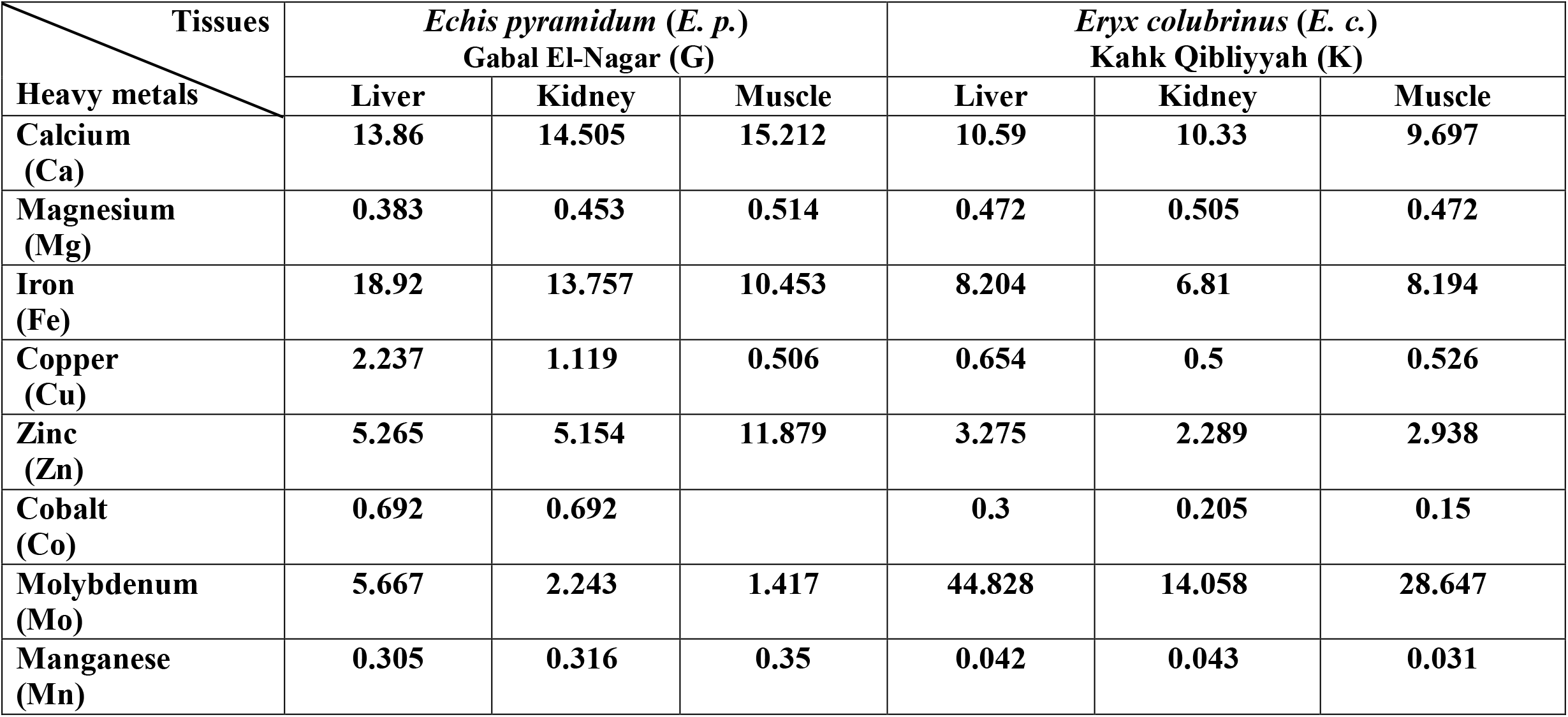
Bioaccumulation factors of essential heavy metals.

Bioaccumulation factor of non-essential heavy metals in Gabal El-Nagar was higher than in Kahk Qibliyyah except B and Al (Table 11).

**Table (11):** Bioaccumulation factors of non-essential heavy metals

| Tissues<br>Heavy Metals | *Echis pyramidum* (E. p.)<br>Gabal El-Nagar (G) |  |  | *Eryx colubrinus* (E. c.)<br>Kahk Qibliyyah (K) |  |  |
| --- | --- | --- | --- | --- | --- | --- |
|  | Liver | Kidney | Muscle | Liver | Kidney | Muscle |
| Boron (B) | 3.411 | 6.643 | 2.135 | 11.875 | 14.127 | 6.634 |
| Aluminium (Al) | 76.001 | 50.339 | 92.809 | 824.357 | 1078.902 | 759.005 |
| Strontium (Sr) | 3.134 | 3.191 | 2.937 | 1.376 | 1.477 | 1.591 |
| Lead (Pb) | 3.096 | 9.017 | 35.038 | 2.477 | 1.153 | 0.85 |
| Nickel (Ni) | 9.15 | 37.298 | 17.008 | 1.351 | 1.806 | 1.351 |
| Cadmium (Cd) | 3.537 | 3.215 | 3.537 | 2.434 | 2.247 | 1.873 |
| Chromium (Cr) | 77.734 | 33.4 | 39.563 | 40 | 29.764 | 39.213 |

Section of liver of the saw-scaled viper, *Echis pyramidum* showed congestion in central vein with intracytoplasmic vacuolization of hepatocyte (Fig 3 A), and it also showed congestion in the portal vein surrounded by fibers and blood sinusoids that considered a sign of fibrosis (Fig 3 B).

Section of liver of the Kenyan sand boa, *Eryx colubrinus* showed severe dilations of central vein surrounded by fibers with severe degeneration in hepatocytes in the surrounding area (Fig 4 A), it showed focal aggregation of melanin-pigmented cells in the hepatic parenchyma (Fig 4 B), and Fig. 4 C was a magnification of Fig. 4 B.

Section of kidney of the saw-scaled viper, *Echis pyramidum* showed hypertrophy and vacuolization in the lining epithelium of distal convoluted tubules (Fig 5 A), it showed focal extravasation of red blood cells between degenerated distal convoluted tubules (Fig 5 B), it also showed hyalinization in the connective tissue stroma between the degenerated distal convoluted tubules (Fig 5 C).

Section of kidney of the Kenyan sand boa, *Eryx colubrinus* showed hypertrophy and vacuolization in the lining epithelium of distal convoluted tubules (Fig 6 A), it showed course granular eosinophilic cytoplasm in the tubular lining epithelium of distal convoluted tubules (Fig 6 B), Fig 6 C was the magnification of Fig 6 B to identify the granular eosinophilic cytoplasm of the lining epithelium of distal convoluted tubules, it also showed pyknosis deep bluish nuclei with deep eosinophilic cytoplasm of some lining tubular epithelium and distal convoluted tubules (Fig 6 D).

Testes of the saw-scaled viper, *Echis pyramidum* showed degeneration in the seminiferous tubules and vacuolization in all the stages of sperm formation associated with the loss of spermiogensis (Fig 7 A), Fig 7 B was magnification of Fig 7 A and Fig 7 C was magnification of Fig 7 B.

**Fig. 6.**
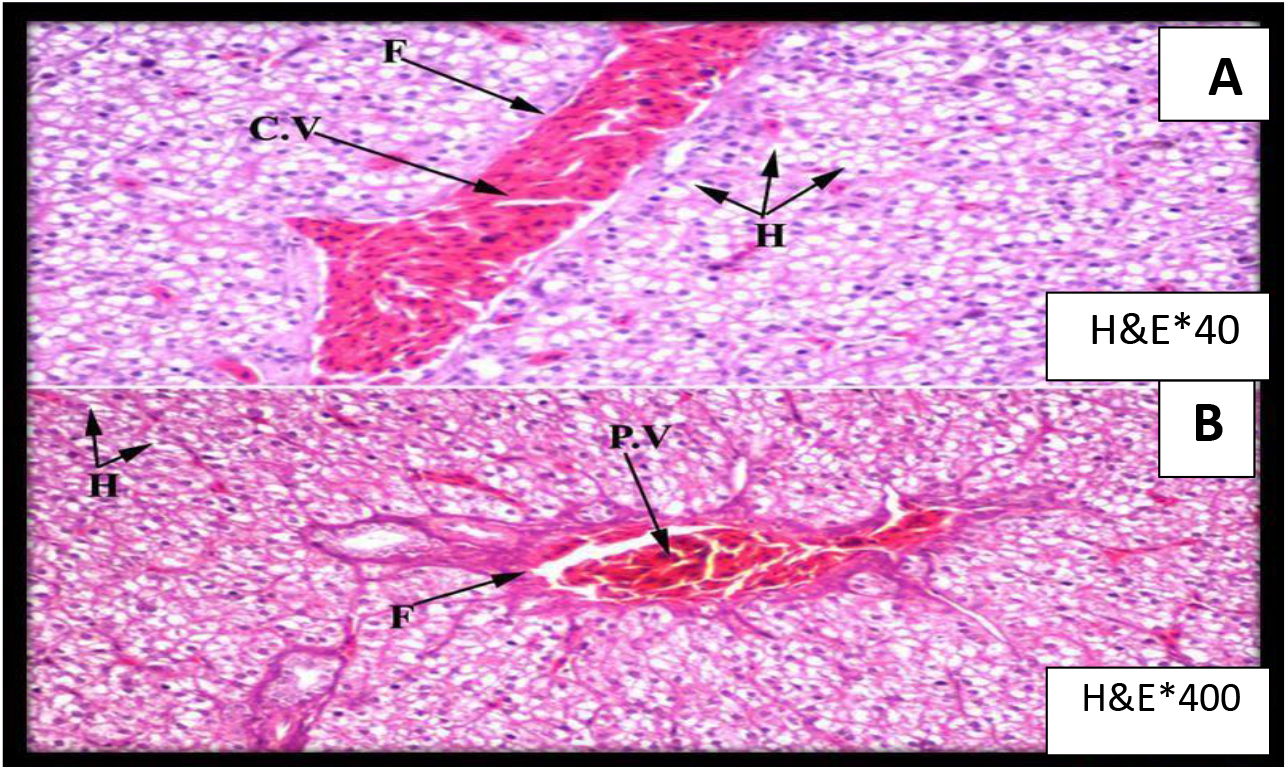
Photomicrograph shows section of liver of saw-scaled viper, *Echis pyramidum*. C.V: Central vein F: Fibers H: Hepatocytes P.V: Portal

**Fig. 7.**
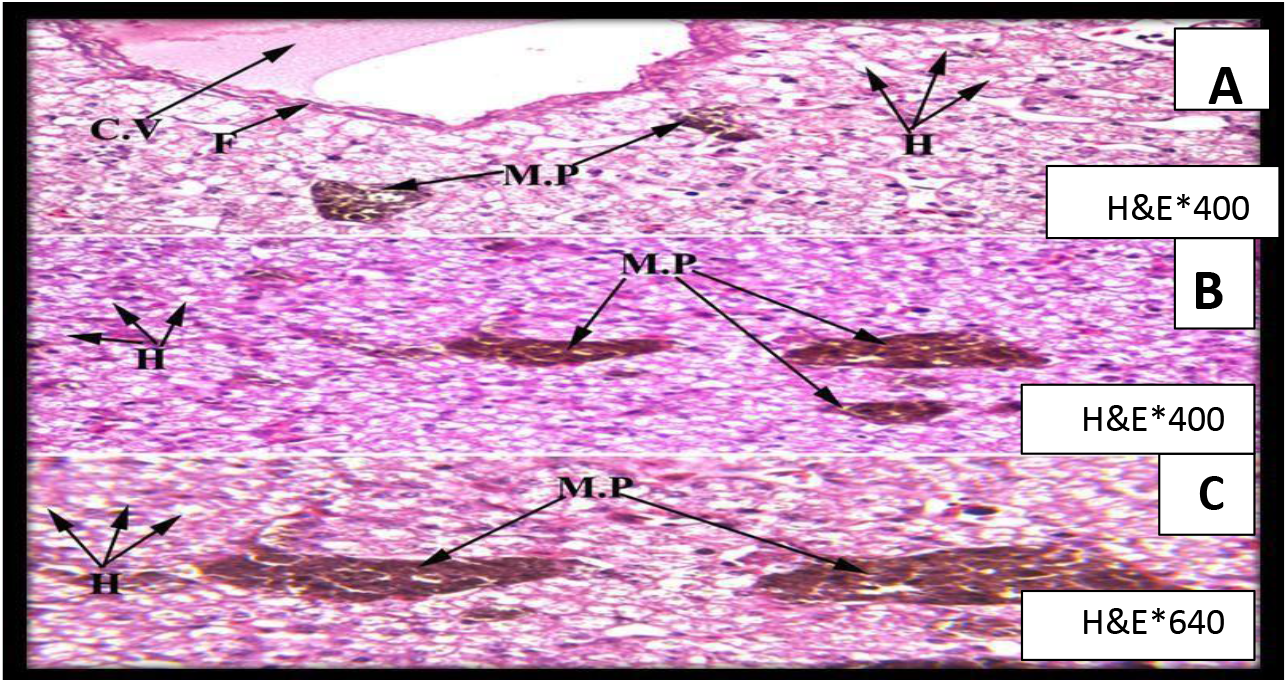
Photomicrograph shows section of liver of Kenyan sand boa, *Eryx colubrinus*. C.V: Central vein F: Fibers H: Hepatocytes M.P: Melanin

Testes of the Kenyan sand boa, *Eryx colubrinus* showed degeneration of some seminiferous tubules (Fig 8 A), Fig 8 B was a magnification of Fig 8 A, and Fig 8 C was a magnification of Fig 8 B.

**Fig. 8.**
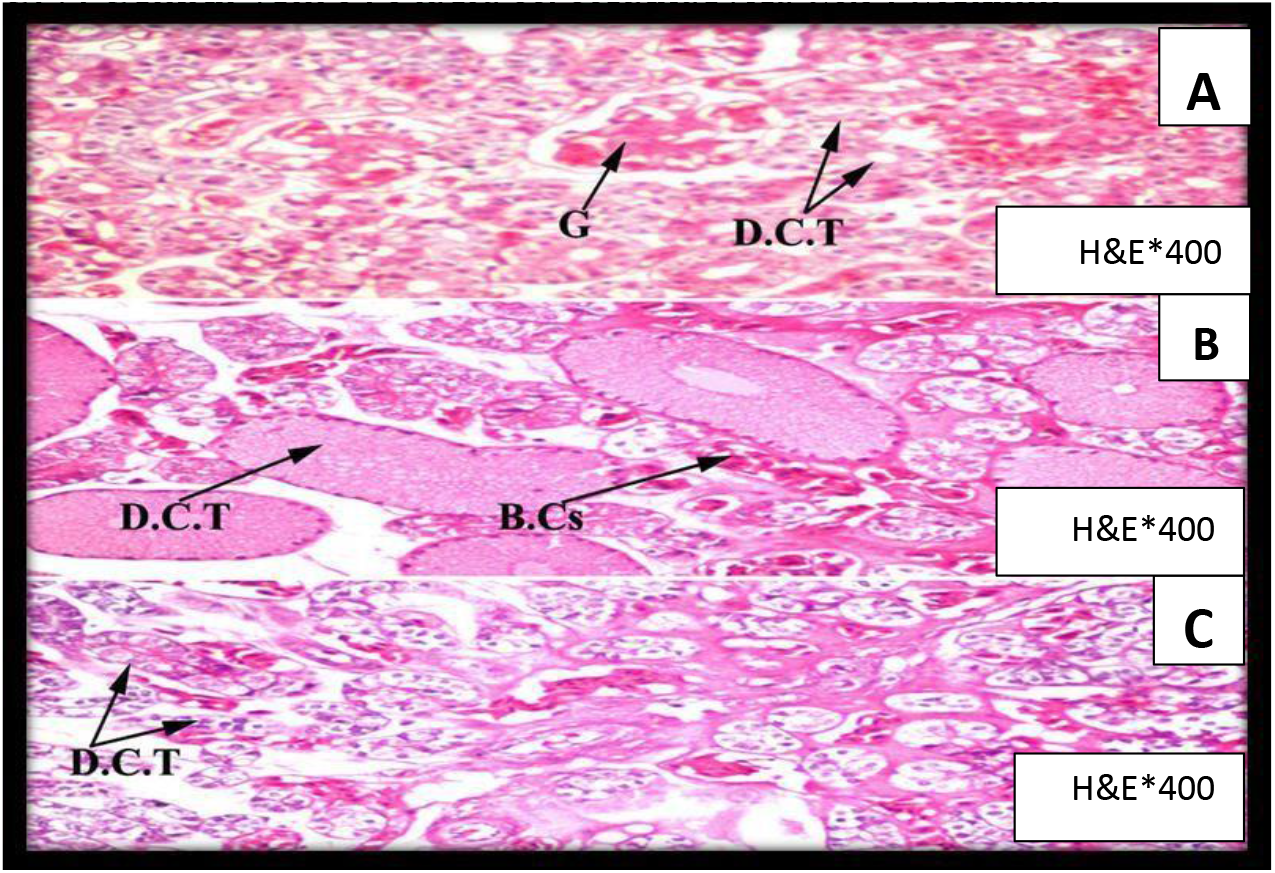
Photomicrograph shows section of kidney of saw-scaled viper, *Echis pyramidum*. G: Glomerulus D.C.T: Distal convoluted tubule B.Cs: Blood cells

**Fig. 9.**
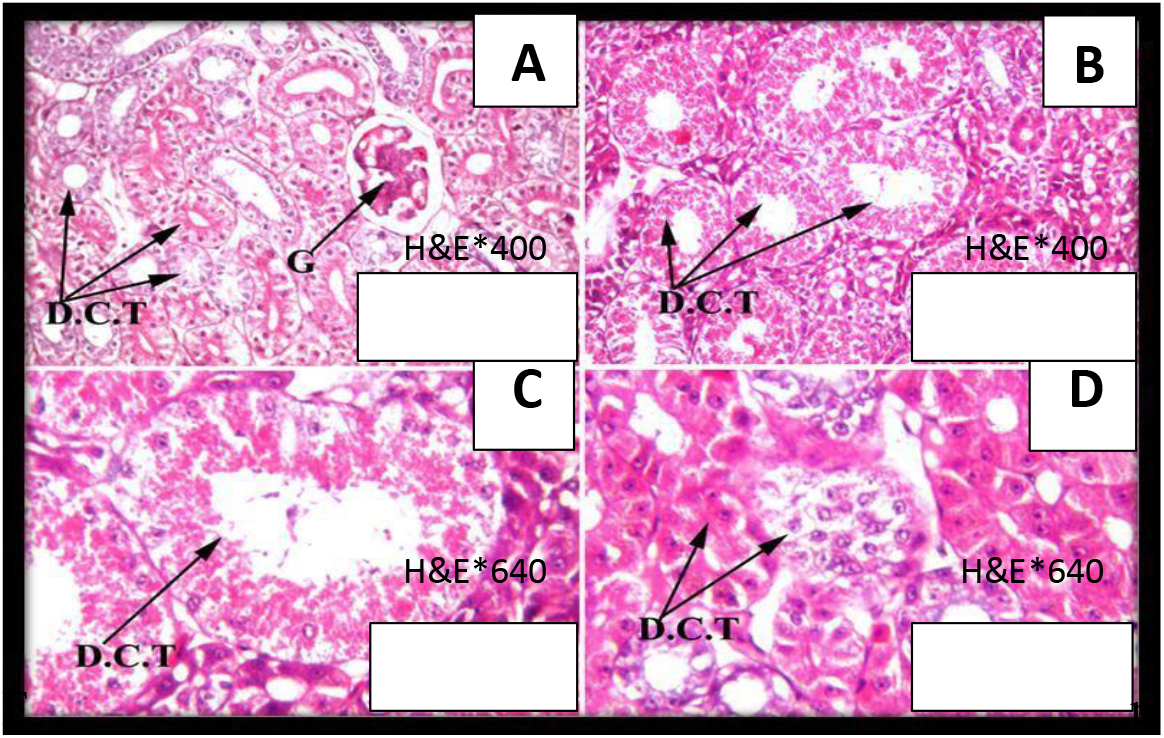
Photomicrograph shows section of kidney of Kenyan sand boa, *Eryx colubrinus*. G: Glomerulus D.C.T: Distal convoluted tubules

**Fig. 10.**
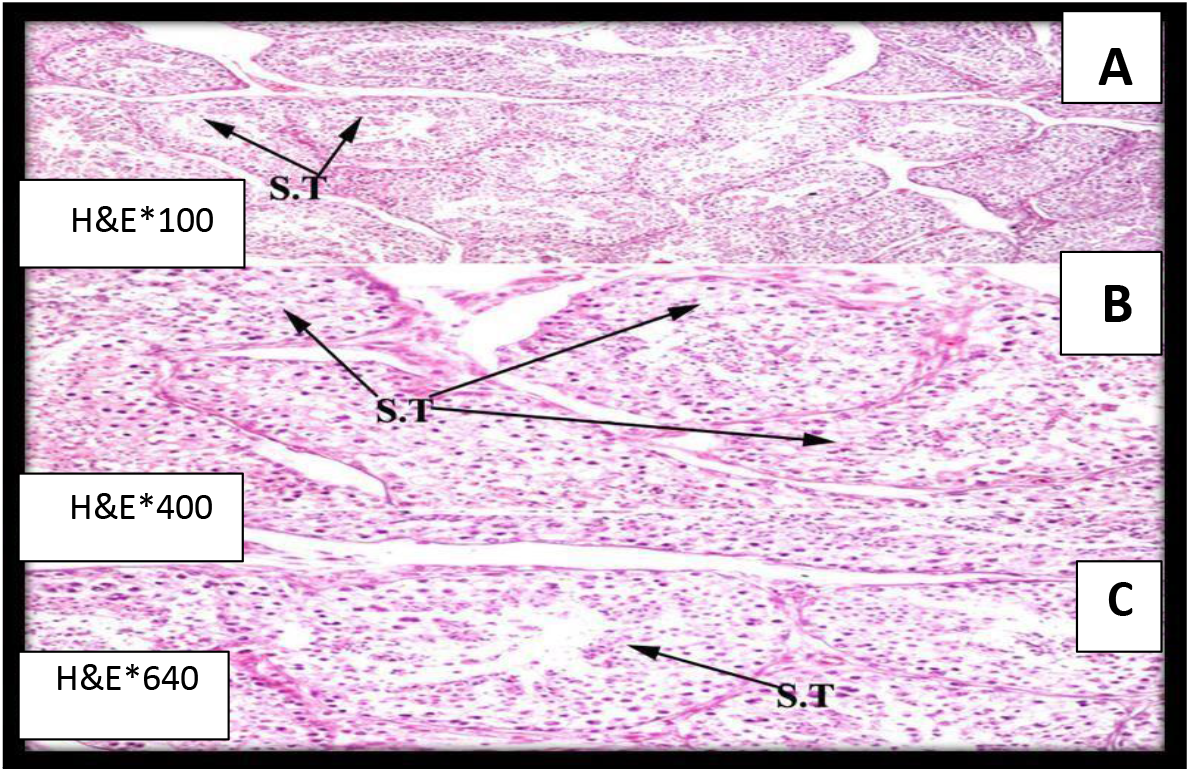
Photomicrograph shows section of testes of saw-scaled viper, *Echis pyramidum*. S.T: Seminiferous tubule

**Fig. 11.**
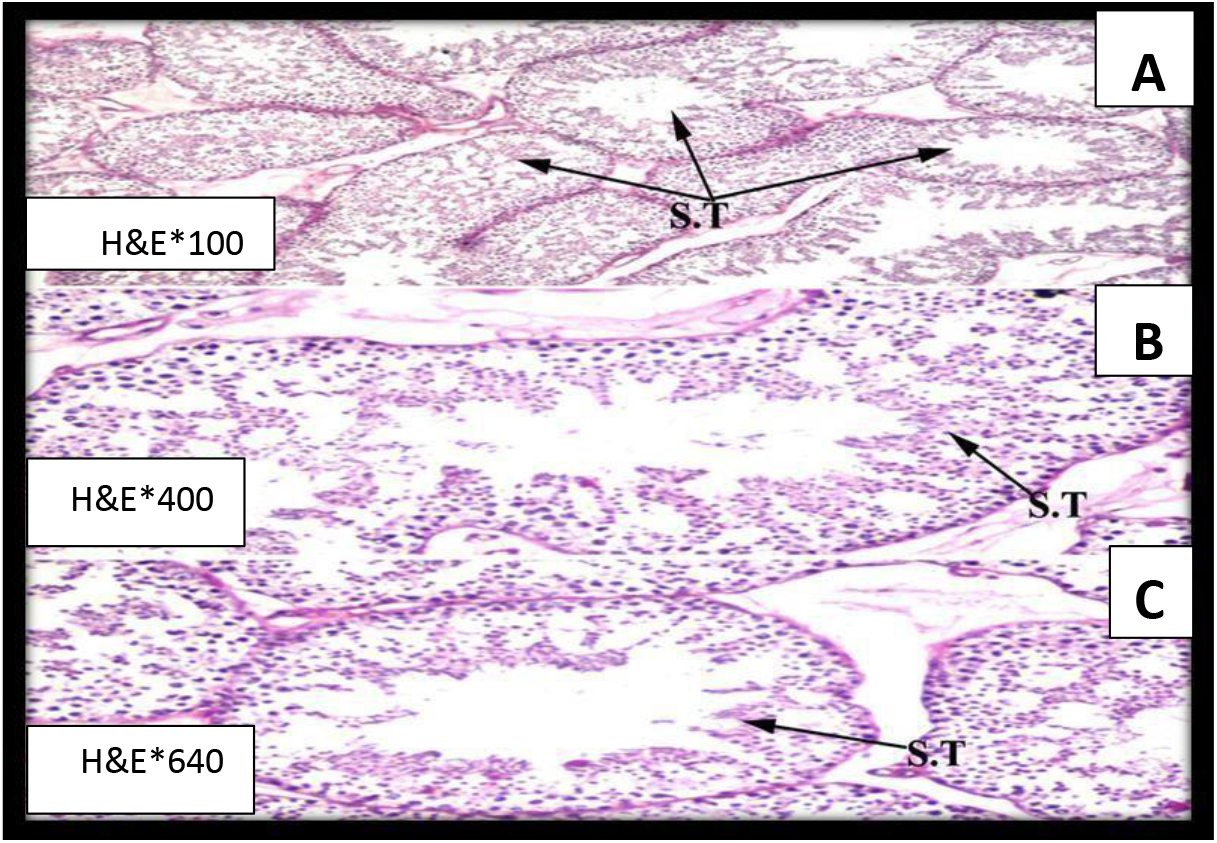
Photomicrograph shows section of testes of Kenayn sand boa, *Eryx colubrinus*. S.T: Seminiferous tubules

## Discussion

Saw-scaled viper, *Echis pyramidum* is a medium sized snake, with a short, stocky body **(El Din, 2006)**. Kenyan sand boa, *Eryx colubrinus* is short and thick snake **(Flower, 1933)**.

The results of the present study indicated an individual variation between the saw-scaled viper, *Echis pyramidum* and the Kenyan sand boa, *Eryx colubrinus* and showed that all the morphometric measurements were higher in the saw-scaled viper, *Echis pyramidum* than in the Kenyan sand boa, *Eryx colubrinus*.

Body weight and diet have influence on the accumulation of metals in snakes **(Drewett *et al.*, 2013)**. Diet is considered the significant route of exposure for the snakes **(Burger *et al.*, 2005)**. Many contaminant studies have reported growth-related changes in metal concentrations in the whole bodies and different tissues of snakes **(Hopkins *et al.*, 1999; Burger *et al.*, 2005; Albrecht *et al.*, 2007 and Rezaie-Atagholipour *et al.*, 2012)**. Growth related changes (body weight, organ weight, hepatosomatic index and gonadosomatic index) in heavy metal concentrations require further investigation in snakes **(Hopkins *et al*., 2005)**.

Results of the present investigation showed a significant increase in the body weight, liver weight, hepatosomatic index, gonads weight, gonadosomatic index, kidney weight and heart weight of the saw-scaled viper, *Echis pyramidum* and significant decrease in the body weight, liver weight, hepatosomatic index, gonad weight, gonadosomatic index, kidney weight and heart weight of the Kenyan sand boa, *Eryx colubrinus*.

Results of the increase in body weight of the saw-scaled viper agreed with **Hopkins *et al* (1999) and Rezaie-Atagholipour *et al*. (2012)** while decrease of the body weight of the Kenyan sand boa agreed with **Burger *et al*. (2005) and Albrecht *et al*. (2007). Hopkins *et al*. (1999) and Rezaie-Atagholipour *et al*. (2012)** found that the increase in the body weight of snakes contaminated with heavy metals was due to bioaccumulation throughout growth.

**Burger *et al*. (2005) and Albrecht *et al*. (2007)** suggested that the decrease in the body weight of snakes contaminated with heavy metals was due to homeostatic control.

Hematology and plasma biochemistry of snakes are considered as useful tools to evaluate the health conditions of individuals especially the adverse effects that are associated with pollution as a result of the exposure to environmental contaminants **(Wilkinson, 2003; Hidalgo-Vila *et al.*, 2007 and Kadry, 2011)**. However, there are no established baseline measurements for a wide variety of blood and plasma biochemical parameters in snakes that can be used as reference values to describe a specific physiological condition **(Gavric *et al.*, 2015)**. Identification of accumulated heavy metals in snakes” tissues provides a measure of the extent of pollutant exposure experienced by organisms but fails to provide insight into the biological significance of exposure to pollutants; therefore, the demand of examining the physiology and/or ecology of snakes in conjunction with tissue concentration of environmental contaminants has been increased **(Hopkins *et al.*, 1999)**.

Results of the present study showed that a significant increase in the blood parameters and calculated blood indices of the saw-scaled viper, *Echis pyramidum* while there is a significant decrease in the blood parameters and calculated blood indices of the Kenyan sand boa, *Eryx colubrinus*.

Results of **Tosunoglu *et al*. (2011)** showed that the venomous snakes exhibited a significant increase in the blood parameters and calculated blood indices, while non-venomous snakes exhibited a significant decrease in the blood parameters and calculated blood indices, and this finding agreed with the present investigation.

The variations in the blood parameters and calculated blood indices could be explained by not only the differences of the normal behavior of each species but also, food and activities of each species **(Vasaruchapong *et al.*, 2013 and Ozzetti *et al.*, 2015)** and they are considered as a useful tool in demonstrating the physiology of snakes after exposure to environmental contaminants **(Wack *et al.*, 2012)**. The difference in blood parameters among species was due to some physiological factors such as sex, age, pregnancy, physical exercise, weather, stress, altitude, captivity and diet **(Alleman *et al.*, 1999; Martínez *et al.*, 2004; Parida *et al.*, 2014 and Gómez *et al.*, 2016)**.

Plasma makes up 60 to 80% of blood volume and contains a great variety of substances in reptiles like other vertebrates present in trace amounts. The concentration of reptilian plasma glucose is associated with seasonal variations. Physiological change within day, especially metabolic cycles is directly correlated with the level of blood glucose in reptiles **(Dessauer, 1970 and Chandavar and Naik, 2004)**. Plasma protein of reptiles comprises 3 to 7% of the total plasma content and it is composed of a mixture of complex protein. Total lipid content of reptilian plasma varies among species according to environmental and physiological conditions **(Dessauer, 1970; Pal *et al.*, 2008 and Parida *et al.*, 2014)**.

Plasma glucose is an important indicator that express the physiological changes in snakes **(Coz-Rakovac *et al.*, 2011)**.

Results of the present study showed a significant increase in the saw-scaled viper, *Echis pyramidum* and a significant decrease in the Kenyan sand boa, *Eryx colubrinus*. In snakes, plasma glucose varies in response to the external environmental fluctuations and reflects their physiological changes as result of their nutrition **(Dessauer, 1970; Campbell, 2004 and Gómez *et al.*, 2016)**. However, different species exhibit different patterns in response to some environmental changes **(Coz-Rakovac *et al.*, 2011)**.

In general, lipid concentrations in snakes seem to be quite conservative and do not change markedly even during prolonged starvation **(Dessauer, 1970 and Coz-Rakovac *et al.*, 2011)**.

Results of the present study showed a significant increase in the saw-scaled viper, *Echis pyramidum* and a significant decrease in the Kenyan sand boa, *Eryx colubrinus*. Elevated plasma lipid could reflect its synthesis in the liver and circulation in blood to other parts of the body for production of fat reserves **(Coz-Rakovac *et al.*, 2011)**. Plasma total protein in snakes is of vital importance in immune and coagulation functions, transportation of ions and hydrophobic molecules, enzymatic activity and regulation of plasma osmolality **(Dessauer, 1970 and Coz-Rakovac *et al.*, 2011)**.

Results of the present study showed a significant increase in the saw-scaled viper, *Echis pyramidum* and a significant decrease in the Kenyan sand boa, *Eryx colubrinus*. Plasma total protein is affected by nutrition **(Dessauer, 1970 and CozRakovac *et al.*, 2011)**. Alternatively, a metabolic disorder, such as insufficient fat stores, may have necessitated unusually high catabolism of protein reserves during hibernation **(Coz-Rakovac *et al.*, 2011)**.

Heavy metals are the most studied class of contaminants in toxicological reports of snakes **(Sparling *et al.*, 2010). Gavrić*et al*. (2015)** mentioned that the position of snakes in food chain makes them very suitable for the accumulation of contaminants and biomonitoring, and also snakes play important ecological roles in controlling the flow of nutrients, energy and contaminants in food webs. Bioaccumulation of metals have been examined in tissues of snakes **(Burger, 1992; Hopkins *et al.*, 1999; Hopkins *et al.*, 2001; Hopkins *et al.*, 2002; Hopkins *et al.*, 2004; Hopkins *et al.*, 2005; Burger *et al*., 2005; Campbell *et al*., 2005; Rainwater *et al*., 2005; Burger *et al.*, 2006; Burger *et al.*, 2007; Wylie *et al.*, 2009; Rezaie-Atagholipour *et al.*, 2012; Drewett *et al.*, 2013 and Serseshk and Bakhtiari, 2015)**. Since the bioaccumulation of metals has been established, the present study focused on the soil as a media transmitting heavy metals into the snakes’ tissues and the biological consequences as a result of heavy metals uptake into the snakes’ tissues.

The relationships between the concentration of heavy metals in snakes and those in soil should be examined from their respective collection sites **(Rainwater *et al*., 2005)**. In this report, the status of essential heavy metals (Ca, Mg, Fe, Cu, Zn, Co, Mo and Mn) and non-essential heavy metals (B, Al, Sr, Pb, Ni, Cd and Cr) in Gabal El-Nagar and Kahk Qibliyah in El-Faiyum, Egypt was studied and associated with the bioavailability of heavy metals in the saw-scaled viper, *Echis pyramidum* and the Kenyan sand boa, *Eryx colubrinus*. El-Faiyum’s soils are alkaline in nature and rich in CaCÜ3 **(Abd Elgawad *et al.*, 2007)**.

The results of our findings revealed that the concentrations of DTPA heavy metals in the surface of soil are variable (Table 7&8), and this agrees with **Abd Elgawad *et al*(2007)** and **Abdel Kawy and Belal (2012)** who state that the surface layer of El-Faiyum’s soils has been subjected to heavy metals as result of atmospheric depositions, applied commercial fertilizers (phosphates in particular), pesticides, manures, waste disposals and may be discharge of untreated domestic sewage and they also suggested that top soils of El-Faiyum have been subjected to heavy metal contaminations and this because of the fact of being close to the main roads and urban areas and also using drainage water as a source of irrigation water in El-Faiyum depression enhance the toxicity of surface soil by increasing heavy metal accumulation because drainage water has high heavy metal content as result of pollution effects, finally these factors jeopardize the health status of the surrounding biota.

There are few reports of the metal levels in terrestrial snakes **(Cocking *et al.*, 1991 and Cristol *et al.*, 2008)**. Variation in habitat preferences (e.g., aquatic vs terrestrial) among snakes makes it possible to evaluate contaminant exposure across habitat types (Campbell and Campbell, 2002). This study examined the accumulation of essential heavy metals (Ca, Mg, Fe, Cu, Zn, Co, Mo and Mn) and non-essential heavy metals (B, Al, Sr, Pb, Ni, Cd and Cr) in the saw-scaled viper, *Echis pyramidum* inhabiting Gabal El-Nagar and the Kenyan sand boa, *Eryx colubrinus* inhabiting Kahk Qiblyiah El-Faiyum.

**Rezaie-Atagholipour *et al*. (2012)** used the prey items of annulated sea snakes to prove that the fact that snakes as predators are able to bio-magnify environmental contaminants within ecosystems and they are also able to transmit contaminants into food chain because they have conversion efficiencies and can convert a large amount of ingested energy into biomass which may be associated with bio-magnification of environmental contaminants.

In this study, we investigate the relationship between the contaminant concentration in the liver, kidney and muscle tissue in the saw-scaled viper and the Kenyan sand boa and their relations with the soils from the collected sites.

The simplest explanation for the difference between our field study and the previous studies conducted by the authors in laboratories **(Hopkins *et al.*, 1999 and Hopkins *et al.*, 2001)** was that snakes collected from field had been exposed to heavy metals for a longer period of time and thus the chance to accumulate heavy metals would be higher. No doubt that the levels of metals within the tissues of snakes are variable **(Burger, 1992 and Burger *et al.*, 2007)**: (1) Ca, Mg, Zn, Mn, Al and Pb were the highest in the muscle of the saw-scaled viper, *Echis pyramidum*, (2) Fe, Cu, Mo and Cr were the highest in the liver of the saw-scaled viper, *Echis pyramidum*, (3) B and Sr were the highest in the kidney of the saw-scaled viper, *Echis pyramidum*, (4) Cd was the highest in the liver and muscle of the saw-scaled viper, *Echis pyramidum*, (5) Co was the highest in the liver and kidney of the saw-scaled viper, *Echis pyramidum*,while the levels of Co in the muscle of the saw-scaled viper, *Echis pyramidum* were not detected. **Nasseem and Abdalla (2003)** observed that pasture soils of the texture of loomy sands and clays, from the North Western Coast of Egypt contain acetic acid soluble EDTA extractable Co from 3.6 to 4.69mg/Kg and it caused Co deficiency in the terrestrial animals inhabiting this soil. They stated that the main factors that cause Co deficiency in terrestrial animals were alkaline and calcareous soils and found that the soils located in the main road mainly faced dust containing polluted Co, which had an influence on the deficiency of Co in the tissues of the terrestrial animals inhabiting these sites. This is interpretation was closely related to the results and properties of the soil of the collected sites, (6) Ca, Fe, Cu, Zn, Co, Mo, Pb, Cd and Cr were the highest in the liver of the Kenyan sand boa, *Eryx colubrinus*, (7) Mg, Mn, B, Al and Ni were the highest in the kidney of the Kenyan sand boa, *Eryx colubrinus*. These findings are associated with **Burger *et al*. (2007)** assumption that not all tissues can be used as indicator of heavy metals exposure. Since Few studies have examined the effect of metal accumulation on snakes **(Campbell and Campbell, 2001)**, and data demonstrating the toxicity threshold in snakes is currently lacking **(Hopkins *et al*., 2005)**, thus; this data in addition to **Burger *et al*. (2007)** data can be used to develop indicators of the relationship among tissue levels, that can be used for predicting levels in other tissues that might not be available.

The levels of the metals accumulated in the liver, kidney and muscle of the saw-scaled viper, *Echis pyramidum* and Kenyan sand boa, *Eryx colubrinus* were variable due to the biogeochemical properties of the collected sites **(Gaines *et al*., 2002; Lord *et al*., 2002; Burger *et al.*, 2006 and Burger *et al.*, 2007)** and the fact that the saw-scaled viper, *Echis pyramidum* belongs to venomous snakes while the Kenyan sand boa, *Eryx colubrinus* belongs to non-venomous snakes, according to this fact their ecology and habitat are different **(El Din, 2006)**, consequently; their methodology of food intake are different. Saw-scaled viper, *Echis pyramidum* track their prey animals, after venom biting in a process called prey relocalization which is done via a protein in their venom **(Hayes *et al.*, 2002)**. Kenyan sand boa, *Eryx colubrinus* track their prey animals by constriction and coil themselves around their prey animals **(Stidworthy, 1974 and Mehrtens, 1987). Burger *et al*. (2007)** stated that different species of snakes are predators and exhibit exposure difference due to differences in prey composition, or to interspecific differences because of toxicodynamics.

Both Saw-scaled viper, *Echis pyramidum* and Kenyan sand boa, *Eryx colubrinus* have the same diet mainly fed on lizards and small mammals such as rodents **(Saleh, 1997)**. According to **El-Din (2006)** *Eryx colubrinus* is a nocturnal snake; partly fossorial, capable of moving below soil surface and waits concealed for its prey. It moves in a distinct serpentine motion above the surface; While Saw-scaled viper, *Echis pyramidum* was found in Faiyum in deeply fissured, dry fluvial soil in the cultivated area. Since the two species of snakes are closely related in feeding behavior. It thus seems that there are interspecific differences in uptake and bioaccumulation among the species that bears examination.

Bioaccumulation factor (BAF) is used to quantify the bioaccumulation of environmental pollutants in aquatic and terrestrial biota **(Mountouris *et al*., 2002)**, and is crucial to estimate chemical residues in biota from measured concentrations in the appropriate reference media **(Hsu *et al.*, 2006)**. Since the ecology of the saw-scaled viper, *Echis pyramidum* and the Kenyan sand boa, *Eryx colubrinus* are different, therefore; the bioaccumulation factors of metals are considered useful indices to estimate the difference between the two species **(Dai *et al*., 2004)**.

Results of our investigation showed that BAF of Mg and Mn in the liver, kidney and muscle of the saw-scaled viper, *Echis pyramidum* and the Kenyan sand boa, *Eryx colubrinus*, BAF of Cu in the muscle of the saw-scaled viper, *Echis pyramidum*, BAF of Cu in the liver, kidney and muscle of the Kenyan sand boa, *Eryx colubrinus*, BAF of Co in the liver and kidney of the saw-scaled viper, *Echis pyramidum* and BAF of Co in the liver, kidney and muscle of the Kenyan sand boa, *Eryx colubrinus* were lower than 1, which means that El-Faiyum’s soil is not contaminated with these metals. While BAF of Ca, Fe, Zn and Mo in the liver, kidney and muscle of the saw-scaled viper, *Echis pyramidum* and the Kenyan sand boa, *Eryx colubrinus* and BAF of Cu in the liver and kidney of the saw-scaled viper, *Echis pyramidum* were higher than 1. BAF of Ca, Fe, Zn and Mo were higher in the saw-scaled viper, *Echis pyramidum* than the Kenyan sand boa, *Eryx colubrinus*, while BAF of Zn was higher in the Kenyan sand boa, *Eryx colubrinus* than the saw-scaled viper, *Echis pyramidum* (Table 10), while BAF of non-essential metals were higher than 1 except BAF of Pb in muscle of the Kenyan sand boa, *Eryx colubrinus* (Table 11).

The variation of metal bioavailability concentrations from soil to snakes was due to the trophic position of snakes within the food web **(Murray, 2003). Hsu *et al*. (2006)** indicated that BAF of metals < 1 was considered a clue that the surface soil was not contaminated with metals, in contrast; BAF of metals > 1 revealed that the tissues of snakes were highly uploaded with metals, particularly toxic metals and it was a proof of the polluted nature of terrestrial food chain.

Toxicity of metals can lead to cell dysfuction, which is considered a sign of necrosis **(Flora *et al*., 2008)**. Metals can either increase or decrease enzymatic activities within the tissues and can lead to histopathological alterations in tissues **(Deore and Wagh, 2012)**. Snakes, unlike the warm-blooded animals, have a poor ability to detoxify toxic heavy metals absorbed and inhaled, or ingested by them with contaminated food because of their low metabolic rates and relatively simple enzyme systems **(Walker and Ronis, 1989)**. This may explain why they are more sensitive to heavy metals so histopathological alterations are induced **(Ganser *et al*., 2003)**.

Liver of the saw-scaled viper, *Echis pyramidum* and the Kenyan sand boa, *Eryx colubrinus* showed liver fibrosis and this agreed with **Ganser *et al*. (2003)** study, which stated that liver fibrosis is the most remarkable pathology in snakes exposed to heavy metals contaminant.

**Schaffner (1998)** stated that chronic toxic injury resulting from environmental pollutants usually leads to liver fibrosis in snakes. Liver of the Kenyan sand boa, *Eryx colubrinus* was more damaged than liver of the saw-scaled viper *Echispyramidum*.

In short, snakes exposed to inorganic contaminants and ingest polluted diet exhibit liver pathology and alteration in liver structure **(Ganser *et al.*, 2003)**. Such abnormalities were conspicuous in nearly one-third of snakes exposed to the contaminated diet, yet exposed snakes otherwise appeared healthy **(Hopkins *et al*., 2002)**.

The kidney of snakes seems to be an important heavy metals metabolizing organ, hence; nephrotoxicity is induced and is considered a target organ of heavy metal’s accumulation **(McClellan-Green *et al*., 2005)**.

In the present study, kidney of saw-scaled viper, *Echis pyramidum* and Kenyan sand boa, *Eryx colubrinus* exhibited hypertrophy and vacuolization of distal convoluted tubules. Kidney of Kenyan sand boa, *Eryx colubrinus* was more damaged than kidney of saw-scaled viper, *Echis pyramidum*. Kidney has been extensively studied in the field of bioaccumulation by **(Hopkins *et al.*, 2001; Hopkins *et al.*, 2002; Hopkins *et al.*, 2004; Hopkins *et al.*, 2005; Burger *et al.*, 2005; Campbell *et al.*, 2005; Rezaie-Atagholipour *et al.*, 2012 and Sereshk and Bakhtiari, 2015)**.

Snakes are considered the best reptilian model for studying reproductive toxicity because they reach reproductive maturity faster than any other reptiles, so they can be better choice for reproductive studies **(Willingham, 2005)**.

Both saw-scaled viper, *Echis pyramidum* and Kenyan sand boa, *Eryx colubrinus* exhibit degeneration of seminiferous tubules. Bioaccumulation of heavy metals in testes has been studied by **(Hopkins *et al.*, 2001; Hopkins *et al.*, 2002; Hopkins *et al.*, 2005 and Burger *et al.*, 2005)**.

## Conclusion

The relationship between the accumulations of heavy metals in different tissues is considered valuable in understanding the ecotoxicology of snakes and enhances our knowledge of the bioavailability cycles of the heavy metals. The demand of understanding the factors that influence heavy metals exposure, bioaccumulation and the consequences of toxic metals on biology and ecology of snakes have been increased. In this report, biological studies and media (soil) through which heavy metals are transmitted to the snakes are investigated to help in the choice of the most suitable bioindicator. According to field observation; saw-scaled viper, *Echis pyramidum* is diverse, while Kenyan sand boa, *Eryx colubrinus* is scarce. Results of the present investigation showed that the saw-scaled viper, *Echis pyramidum* can withstand hard conditions more than the Kenyan sand boa, *Eryx colubrinus*. Therefore, Kenyan sand boa, *Eryx colubrinus* is considered the most suitable bioindictor monitoring heavy metals as environmental contaminants.

In Egypt, both saw-scaled viper, *Echis pyramidum* and Kenyan sand boa, *Eryx colubrinus* were classified as Least Concern, but as a measure of their conservation status in their contaminated habitat, a Red Spot in the IUCN Red List of saw-scaled viper, *Echis pyramidum* and Kenyan sand boa, *Eryx colubrinus* was suggested to categorized them as Near Threatened and Vulnearable, respectively (IUCN, 2005).

